# Tilorone: A Broad-Spectrum Antiviral For Emerging Viruses

**DOI:** 10.1101/2020.03.09.984856

**Authors:** Sean Ekins, Peter B. Madrid

## Abstract

Tilorone is a 50-year-old synthetic small-molecule compound with antiviral activity that is proposed to induce interferon after oral administration. This drug is used as a broad-spectrum antiviral in several countries of the Russian Federation. We have recently described activity *in vitro* and *in vivo* against the Ebola Virus. After a broad screening of additional viruses, we now describe *in vitro* activity against Chikungunya virus (CHIK) and Middle Eastern Respiratory Syndrome Coronavirus (MERS-CoV).

---

In recent years we have witnessed major Ebola virus outbreaks in Africa, the Zika outbreak in Brazil and now the novel coronavirus (2019-nCoV) in China. These viruses are rarely confined to their original locations and thus create challenges in containment. Newer viruses also lack available approved treatments and indicate the need for broader spectrum antivirals. We now highlight one such molecule, tilorone dihydrochloride (tilorone, Amixin^®^ or Lavomax^®^) which is currently registered for human use in Russia, Ukraine, Kazakhstan, Belarus, Armenia, Georgia, Kyrgyzstan, Moldova, Turkmenistan, and Uzbekistan as an antiviral (influenza, acute respiratory viral infection, viral hepatitis, viral encephalitis, myelitis, and others) and immunomodulating medication. It is also included in the list of essential medicines of the Russian Federation. *In vivo* efficacy studies dating back to 1970 support possible uses against a broad array of viruses including influenza A, influenza B, herpes simplex virus 1, West Nile virus, Mengo virus, Semliki Forest virus, vesicular stomatitis virus and encephalomyocarditis virus (1–3). More recently we recently demonstrated 90-100% survival in mice infected with Ebola then treated with tilorone (4). These results led us to more broadly profile the antiviral spectrum of activity and focus on Chikungunya virus (CHIK) and Middle Eastern Respiratory Syndrome Coronavirus (MERS-CoV).

## Tilorone screening in antiviral assays

Tilorone dihydrochloride was purchased from Sigma-Aldrich (St. Louis, MO). Tilorone was tested (using the NIAID DMID services) against representatives of the herpesviridae, bunyaviridae, togaviridae, arenaviridae, flavivirdae, picornaviridae, poxviridae, hepatic viruses, respiratory viruses and other viruses. Four-concentration CPE inhibition assays were performed. Confluent or nearconfluent cell culture monolayers in 96-well disposable microplates were prepared. Cells were maintained in MEM or DMEM supplemented with FBS as required for each cell line. For antiviral assays the same medium was used but with FBS reduced to 2% or less and supplemented with 50 μg/ml gentamicin. The test compound is prepared at four log_10_ final concentrations 0.1, 1.0, 10, and 100 μg/ml or μM. Five microwells are used per dilution: three for infected cultures and two for uninfected toxicity cultures. Controls for the experiment consist of six microwells that are infected (virus controls) and six that are untreated (cell controls). The virus control and cell control wells are on every microplate. In parallel, a known active drug is tested as a positive control drug using the same method as is applied for test compounds. The positive control is tested with each test run. The assay was initiated by first removing growth media from the 96-well plates of cells. Then the test compound was applied in 0.1 ml volume to wells at 2X concentration. Virus, normally at <100 50% cell culture infectious doses (CCID_50_) in 0.1 ml volume, was placed in those wells designated for virus infection. Medium devoid of virus was placed in toxicity control wells and cell control wells. Virus control wells were treated similarly with virus. Plates are incubated at 37°C with 5% CO_2_ until maximum CPE is observed microscopically in virus control wells. The plates are then stained with 0.011% neutral red for approximately two hours at 37°C in a 5% CO_2_ incubator. The neutral red medium was removed by complete aspiration, and the cells rinsed 1X with phosphate buffered solution (PBS) to remove residual dye. PBS was completely removed and the incorporated neutral red eluted with 50% Sorensen’s citrate buffer/50% ethanol for at least 30 minutes. The dye content in each well was quantified using a 96-well spectrophotometer at 540 nm wavelength. The 50% effective (EC_50_, virus-inhibitory) concentrations and 50% cytotoxic (CC_50_, cell-inhibitory) concentrations were then calculated by linear regression analysis. The quotient of CC_50_ divided by EC_50_ gives the selectivity index (SI_50_) value. Ideally compounds showing SI_50_ values >10 are considered active.

## Tilorone has in vitro activity against CHIK and MERS

We identified promising micromolar activities for Tilorone against CHIK and MERS-CoV with reasonable selectivity indexes (Table 1). The *in vitro* activity against MERS also agrees with recent findings of others (5). These combined observations along with earlier descriptions of many antiviral activities suggest tilorone is a potential broad-spectrum antiviral that may have utility against additional coronaviruses. While this drug is approved in Russia Federation countries, Tilorone has never been evaluated and tested for safety and efficacy under studies that meet current ICH and FDA guidelines and regulations. Recent virus outbreaks such as SARS-CoV-2 suggest the urgent need for reassessment of this compound as a broad-spectrum antiviral as we have yet to fully appreciate the utility of this drug discovered 50 years ago.

**Table 1.** Antiviral screening data for Tilorone generated under the NIAID-DMID NCEA antiviral *in vitro* screening services. * *In vitro* antiviral data in Vero 76 cells may underestimate antiviral activity due to lacking IFN pathways.

| Virus | Strain | Genus | Type | Cell line | EC<sub>50</sub><br>(μM) | CC<sub>50</sub><br>(μM) | SI |
| --- | --- | --- | --- | --- | --- | --- | --- |
| CHIKV | S27 (VR-67) | Alphavirus | +<br>ssRNA | Vero 76 | 4.2* | 32 | 7.6 |
| MERS-CoV | EMC | Betacoronavirus | +<br>ssRNA | Vero 76 | 3.7* | 36 | 9.7 |

## ACKNOWLEDGMENTS

Dr. Mindy Davis is gratefully acknowledged for assistance with the NIAID virus screening capabilities, Task Order number B22.

